# Identification of a unique transcriptomic signature associated with islet amyloidosis

**DOI:** 10.1101/2022.11.09.515784

**Authors:** Marko Barovic, Klaus Steinmeyer, Nicole Kipke, Eyke Schöniger, Daniela Friedland, Flavia Marzetta, Almuth Forberger, Gustavo Baretton, Jürgen Weitz, Daniela Aust, Mark Ibberson, Marius Distler, Anke M Schulte, Michele Solimena

**Author notes:** current contact address (Anke M Schulte).

## Abstract

**Aims:** This cross-sectional study aims to identify potential transcriptomic changes conveyed by presence of amyloid deposits in islets from pancreatic tissue obtained from metabolically profiled living donors.

**Methods:** After establishing Thioflavin S as the most sensitive approach to detect islet amyloid plaques, we utilized RNA sequencing data obtained from laser capture microdissected islets to define transcriptomic effects of this pathological entity. The RNA sequencing data was used to identify differentially expressed genes by linear modeling. Further analyses included functional enrichment analysis of KEGG and Hallmark gene sets as well as a weighted gene correlation network analysis.

**Results:** Eleven differentially expressed genes were identified in islets affected by amyloidosis. Enrichment analyses pointed to signatures related to protein aggregation diseases, energy metabolism and inflammatory response. A gene co-expression module was identified that correlated to islet amyloidosis.

**Conclusion:** Although the influence of underlying Type 2 diabetes could not be entirely excluded, this study presents a valuable insight into the biology of islet amyloidosis, particularly providing hints into the potential relationship between islet amyloid deposition and structural and functional proteins involved in insulin secretion.

**Research in context:** *What is already known about this subject?:*

- Islet amyloidosis is the only histological marker of Type 2 diabetes in the pancreas
- Individuals not suffering from Type 2 diabetes can also be affected by islet amyloidosis
- The clinicopathological significance of this phenomenon is still unclear

*What is the key question?:*

- Does the islet transcriptome of individuals with islet amyloidosis provide explanations for the onset of this phenomenon and its pathophysiological value?

*What are the new findings?:*

- Islet transcriptomes of affected subjects exhibit only limited transcriptomic differences compared to unaffected ones.
- Structural and functional proteins involved in insulin secretion machinery may be involved in the pathophysiological sequence of amyloid formation

## Introduction

Type 2 diabetes mellitus (T2D) is a metabolic disorder characterized by chronic hyperglycemia in the absence of immune-mediated beta cell destruction. The main disease burden is defined through its numerous complications that occur over several years, and negatively affects healthcare systems across the world. This disease is principally characterized by two pathological features: beta cell failure and the resistance of peripheral tissues to the action of insulin. The body of research focusing on beta cell failure as the *conditio sine qua non* for the onset of T2D spans across decades and scientific fields, seeking the mechanisms causing the demise of the beta cell secretory machinery and thus the decompensation of glucose homeostasis.

Diagnostic tests from peripheral blood have been validated and well established in the clinical practice. These provide valuable, easily accessible and therapeutically crucially important information on the metabolic state of the patient. On the other hand, morphological changes in the islets of Langerhans and more specifically the insulin-producing beta cells which could potentially inform research advances in the field of beta cell biology are yet to be discovered, with a notable exception of islet amyloidosis. Hyaline deposits in islets of Langerhans have been identified in human pancreatic tissue sections essentially since the dawn of modern histopathology.(1) This phenomenon was, however, not viewed as particularly significant because of its occurrence also in the tissue of normoglycemic subjects, although in a smaller extent compared to patients with T2D.(2)

Despite the noted inter-species conservation, the minute differences in the peptide sequence of IAPP across species proved to be sufficient to convey a disposition of the human and some other vertebrate variants to fibrillate and form aggregates. These, in turn, are considered to be the precursors of islet amyloid plaques, by serving as deposition nuclei. It remains disputed whether these nuclei come to be in the intercellular space or within the cell, which would explain the cytotoxic effect of amyloid plaques on beta cells given that the extracellular amyloid deposits are considered to be inert. The most recent understanding of the chain of events proposes that the fibrillation indeed starts within the beta cell leading to cell death and release of the seeding aggregates into the intercellular space where they grow by further deposition to form the extracellular plaques nowadays most commonly understood as islet amyloidosis.(3) Notably, the IAPP of the most frequently used experimental animals in diabetes research – mouse and rat – does not have the sequence motifs that confer amyloidogenic properties. For this reason, several transgenic model organisms have been established in order to study islet amyloidosis on a molecular level, each with their own shortcomings in recapitulating this phenomenon in humans.

In this study, we profiled the pancreatic tissue biobank of the LIving DOnor PAncreatic COhort (LIDOPACO) initiative for the presence of islet amyloidosis after establishing a sensitive methodology for its detection by microscopy. This data was then integrated with the extensive clinical data for each individual donor, thereby enabling the establishment of a database on human islet amyloidosis with unprecedented detail. Molecular patterns associated with islet amyloidosis were pursued by means of the analysis of islet RNA sequencing data available for all the donors in this cohort.

## Material and methods

### Cohort

Based on tissue availability, a total of 121 patients enrolled in the LIDOPACO program were included in this study in accordance with the approval of the Ethical Committee of the Technische Universität Dresden. After obtaining informed consent, medical history and clinical laboratory data relevant for metabolic status was gathered either from existing preoperative preparatory examinations or from peripheral blood samples collected with the goal of completing the metabolic profiling as part of the LIDOPACO (e.g. oral glucose tolerance test (OGTT)). According to this data, patients were assigned either to the non-diabetic (ND), impaired glucose tolerance (IGT), pancreoprive diabetes (T3cD) or T2D categories according to the criteria defined by the American Diabetes Association (ADA). The ND group included 18 patients, the IGT group 39 patients classified as pre-diabetic (including patients with impaired fasting glucose) and the T3cD and T2D groups 32 patients each. Patients who underwent neoadjuvant chemotherapy, as well as those with pancreatic neoplasms of endocrine origin were excluded from the study.

### Pancreatic tissue procurement

Immediately following the surgical procedures of partial or total pancreatectomy conducted for various indications at the Clinic for Visceral, Thoracic and Vascular Surgery of the University Hospital Carl Gustav Carus, surgical pathological samples were collected and processed by a pathologist. Fragments of healthy tissue were prepared and either snap frozen in liquid nitrogen and thereafter stored at -80° or fixed in formalin and embedded in paraffin.

### Detection of islet amyloid plaques

Islet amyloid plaques were detected in 2 μm thin sections of formalin fixed paraffin embedded (FFPE) tissue using histochemical labeling by either Congo Red or Thioflavin S. Slides were stained with Congo Red according to the standard protocols of the Institute for Pathology of the University Hospital Carl Gustav Carus and evaluated for presence of amyloidosis in a binary fashion (present/not present) by a pathologist.

Staining with Thioflavin S was performed by incubating the slides for 10 minutes in the staining solution (0.5% Thioflavin S w/v in distilled water) followed by washing steps. Immediately afterwards, the slides were subjected to an automated immunofluorescence protocol on the Roche Ventana Discovery system targeting synaptophysin (Rabbit anti-Synaptophysin (ThermoFisher, MA5-14532, final dilution 1:30); goat anti-rabbit AF750 (Invitrogen, A21039; final dilution 1:150)) without signal amplification and with DAPI as nuclear counterstain. Whole slide images were acquired on a Zeiss AxioScan slide scanner at 20x magnification in the appropriate fluorescence channels.

Across all slide scans, at least 200 islets were manually segmented per sample based on the synaptophysin expression, with the examiner blinded for the Thioflavin S signal (in total 28344 islets). These were saved as regions of interest (ROIs) and used as delimiter for binary detection of a signal in the Thioflavin S channel (present/not present) in each ROI. Accounting for lack of specificity, we set the threshold for classification into the amyloid positive group to be >1 rather than >0 affected islets. Considering that the aimed for number of assessed islets per donor was 200, this corresponds to a threshold of 0.5%, which is widely regarded in statistics as an acceptable error margin. Substantiating our approach of assessing only a small portion of all islets in the whole gland for practicality reasons is the previous finding of homogeneous distribution of amyloidosis affected islets in transgenic mice.(4) All image analysis was done using Fiji version 1.53.

### Transcriptomics

#### Laser Capture Microdissection

Pancreatic islets for RNA sequencing were collected using Laser Capture Microdissection (LCM) as previously described.(5–7) Briefly, snap frozen tissue was sectioned in a cryostat into 20μm thin sections and mounted on Zeiss PEN Membrane Slides. After drying and in strictly RNAse-free conditions, these slides were processed on a Zeiss Palm MicroBeam system. A total islet area of 1×10^6^ μm^2^ was excised based on the islet lipofuscin autofluorescence for each patient. Complete mRNA was isolated using Arcturus PicoPure RNA isolation kit and its concentration was determined using Qubit RNA high sensitivity platform. Before proceeding to the next steps, RNA integrity was determined using an Agilent Bioanalyzer 2100.

#### RNA sequencing

Complementary DNA (cDNA) libraries were constructed using the Illumina Smart-seq2 kit selecting only for the Poly(A)^+^ strands. After enrichment, purification and fragmentation steps, the libraries were subjected to single ended sequencing at the DRESDEN-Concept Genome Center on the Illumina NovaSeq 500 platform with 76bp read length and target depth of at least 35 million reads per library.

Low quality sequences and adapters were filtered using Cutadapt (ver. 1.8), and fastq_screen (ver. 0.11.1) was used for removal of ribosomal sequences. Low complexity reads were subsequently removed with reaper (ver. 15-065) and the remaining reads aligned to the Genome Reference Consortium Human Build 38 patch release 12 index using STAR aligner (ver. 2.5.3a). Read counts per gene locus were summarized using htseq-count (ver. 0.9.1).

Samples were grossly assessed for presence of exocrine contamination by examining the expression levels of genes typical for the exocrine tissue (*PLIP, PRSS1, AMY2A, AMY2B*). This and further quality assurance steps (analysis of the absolute number of detected genes, gene body coverage and cumulative gene diversity) were used for a more granular quality assurance. For downstream analyses, the dataset was filtered for minimal expression retaining only the genes with at least 1 count per million reads (CPM) in at least 10% of the samples.

#### Differential expression and enrichment analysis

Bioconductor package DESeq2 functions *calcNormFactors* and *vst* were used for library size normalization and variance stabilizing transformation respectively. Exploratory multidimensional scaling plots were created from this data using the R function plotMDS (limma). Analysis of differential gene expression was performed using the bioconductor package limma. In an attempt to isolate the effects of islet amyloidosis from the effects of the disease itself, diabetes status was included as covariate in the linear model, together with age, gender, BMI and sequencing batch. Genes with Benjamini-Hochberg multiple comparison corrected *p* value lower than 0.05 and absolute fold change higher than 1.2 were considered differentially expressed.

Functional enrichment analysis of Kyoto Encyclopedia of Genes and Genomes (KEGG) pathways was conducted using Gene Set Enrichment Analysis using the *gseKEGG* function of the R package clusterProfiler (ver. 3.10.1) on unfiltered gene lists ranked by decreasing differential expression adjusted *p* value in an unsigned manner. Gene sets were filtered to include only those sets which contained >100 and <500 genes, and the enrichment scores were normalized by gene set size. Results were filtered using a cutoff of 0.01 on the BH-adjusted *p* value and term redundancy was reduced using the function *simplify* from the same package and using enrichment maps generated by the *emapplot* function from the R package enrichplot (ver. 1.2.0).

Enrichment of The Broad Institute Molecular Signatures Database v7.4 hallmark gene sets was assessed by competitive gene set test accounting for the inter-gene correlation, using the function *camera* from the limma Bioconductor package.

#### Weighted Gene Correlation Network Analysis

The Weighted Gene Correlation Network Analysis (WGCNA) was performed using the R package WGCNA (ver. 1.68). A gene coexpression network was constructed from normalized, batch-corrected (*removeBatchEffect* function from the package limma) and vst transformed expression counts. An unsigned adjacency matrix was constructed using Pearson correlation of pairwise complete observations. Soft threshold parameter was optimized using the *pickSoftThreshold* function and the best threshold (α=6) was visually determined following developers’ recommendations. A topological overlap matrix was derived from the adjacency matrix and converted to distances, which were hierarchically clustered using average linkage clustering. Finally, dynamic tree cut using hybrid method and parameters minClusterSize=20 and deepSplit=2 identified individual gene coexpression modules. Similar modules were in turn merged based on an eigengene distance threshold of 0.15.

Spearman correlation was used to relate the module eigengenes to selected clinical traits and to the qualitative islet amyloidosis status based on the Thioflavin S staining. A heatmap of the module-trait correlations was produced using *labeledHeatmap* function from the WGCNA package, depicting the Spearman correlation coefficients and corresponding *p* values calculated using *cor* and *corPvalueStudent* functions from the same package. In the module with the strongest correlation to islet amyloidosis, genes with the highest correlation to the trait and the module eigengene were selected as module hub genes.

## Results

The cohort used in this study consisted of 121 surgical pancreatic tissue donors participating in the LIDOPACO programme. The patients were included into this study based on the availability of tissue for the necessary investigations. Eighteen of the patients were classified as ND through the preoperative metabolic profiling procedure, 39 were in the IGT group while the T3cD and T2D groups consisted of 32 subjects each (Fig. 1A). In terms of basic demographic/clinical measures, the ND subjects were found to have a significantly lower age and BMI. Expectedly, these subjects also exhibited a lower HbA1c and glycaemia at 2h point of OGTT values compared to the other groups (Fig. 1B). Malignant tumors were markedly more common in the T2D and T3cD groups than in the IGT group, with an even bigger gap when compared to the ND group (Table 1).

**Table 1.** Underlying pancreatic disease. Frequencies of underlying pancreatic disorders categorized by diabetes status

| Diabetes status | Chronic pancreatitis | Benign tumor | Malignant tumor |
| --- | --- | --- | --- |
| ND | 4 (22.22%) | 5 (27.78%) | 9 (50%) |
| IGT | 6 (15.38%) | 9 (23.07%) | 24 (61.53%) |
| T3cD | 5 (15.62%) | 3 (9.37%) | 24 (75%) |
| T2D | 6 (18.75%) | 3 (9.37%) | 23 (71.87%) |

**Figure 1.**
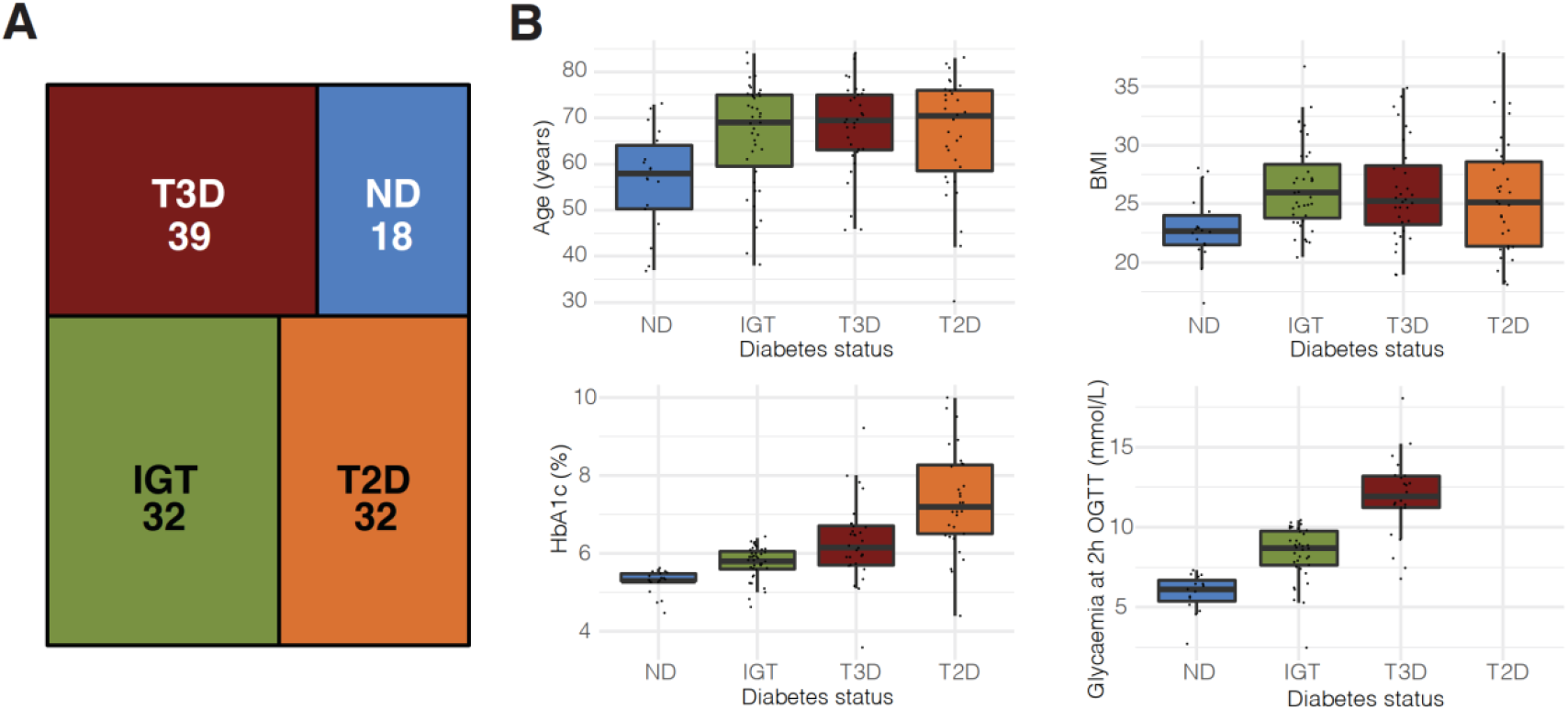
Cohort characteristics overview. **(A)** Treemap plot of the cohort composition in view of diabetes status. **(B)** Basic clinical characteristics of the cohort, separated into diabetes status categories. Boxplot corresponds to the span between the 25^th^ and 75^th^ percentile, with horizontal line marking the median. The whiskers reach to the most extreme data point that is no more than 1.5 times the length of the box away from the box. ND non-diabetic, IGT impaired glucose tolerance, T3cD Type 3c diabetes, T2D Type 2 diabetes.

At first, we aimed to determine the most sensitive method of amyloid plaque detection. To this end, we employed two historically well-established methods for the detection of amyloid plaques – Congo Red and Thioflavin S. The Congo Red staining evaluated by a pathologist revealed only 6 amyloid positive samples. At odds with this finding, a custom semi-automated pipeline applied on the manually segmented fluorescence microphotographs of 28 344 pancreatic islets stained with Thioflavin S, identified islet amyloidosis in 5/18 (27.8%) ND subjects, 16/39 (41%) in IGT, 14/32 (43.7%) in T3cD and 25/32 (78.1%) in T2D subjects (Figure 2A). The islet amyloidosis assignment based on this method was therefore used for all downstream analyses. Representative images of amyloid positive and amyloid negative islets are shown in Figure 2B.

**Figure 2.**
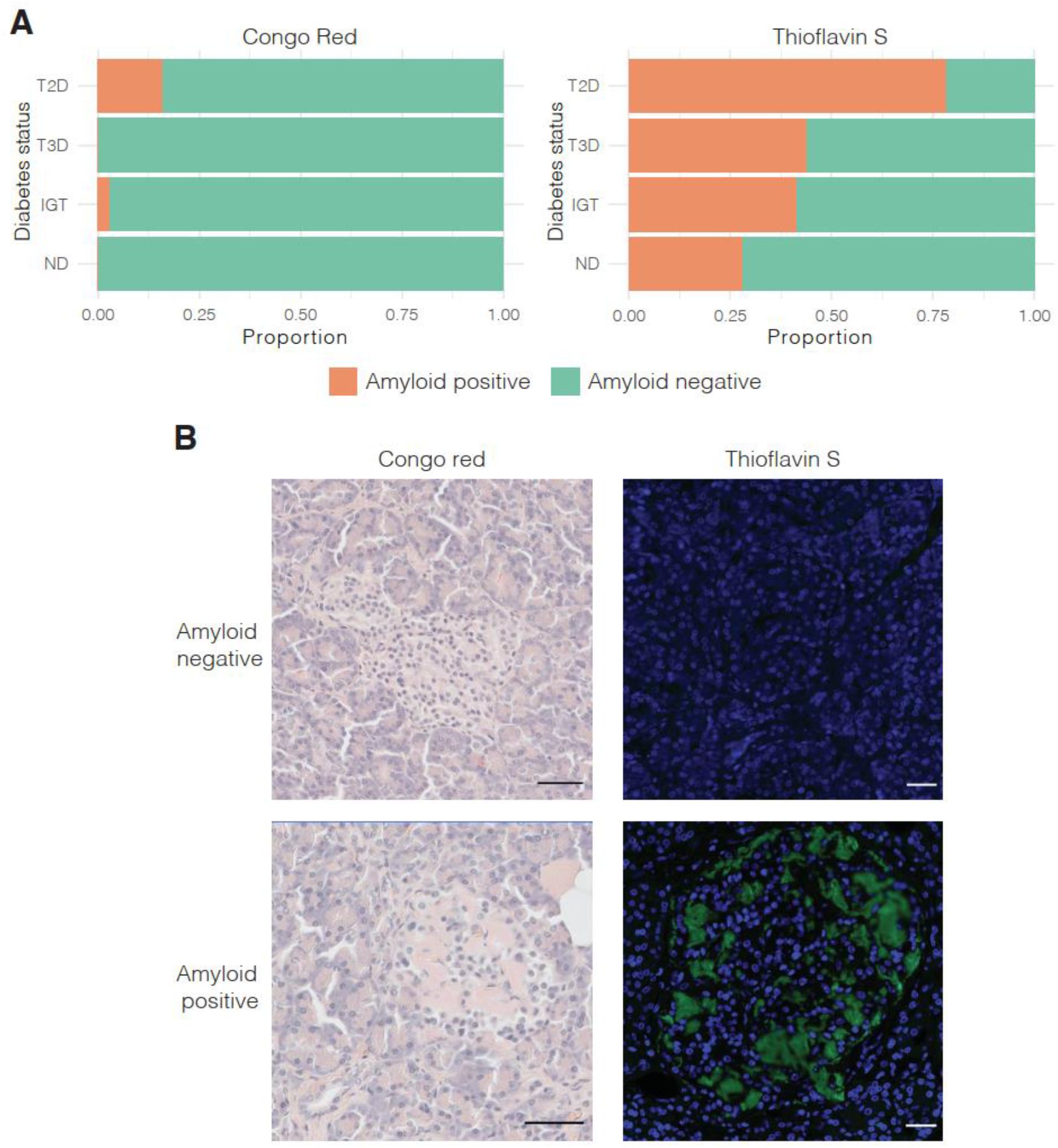
Islet amyloidosis detection. **(A)** Stacked bar chart of proportions between amyloid positive and amyloid negative samples between the two amyloid staining approaches. **(B)** Representative images of Congo red and Thioflavin S staining techniques with positive and negative staining examples. Scale bar corresponds to 50 μm.

Assessment of the basic metabolically relevant clinical variables in the subjects with and without islet amyloidosis unsurprisingly revealed a significantly higher HbA1c levels (Figure 3A). This is consistent with the growing percentage of amyloidosis affected subjects in the disease categories progressing towards overt T2D. Interestingly, no difference was found in age, a demographic parameter typically associated with onset of amyloidosis in general. Same was true for BMI and glycaemia at 2h point of OGTT (Figure 3A).

**Figure 3.**
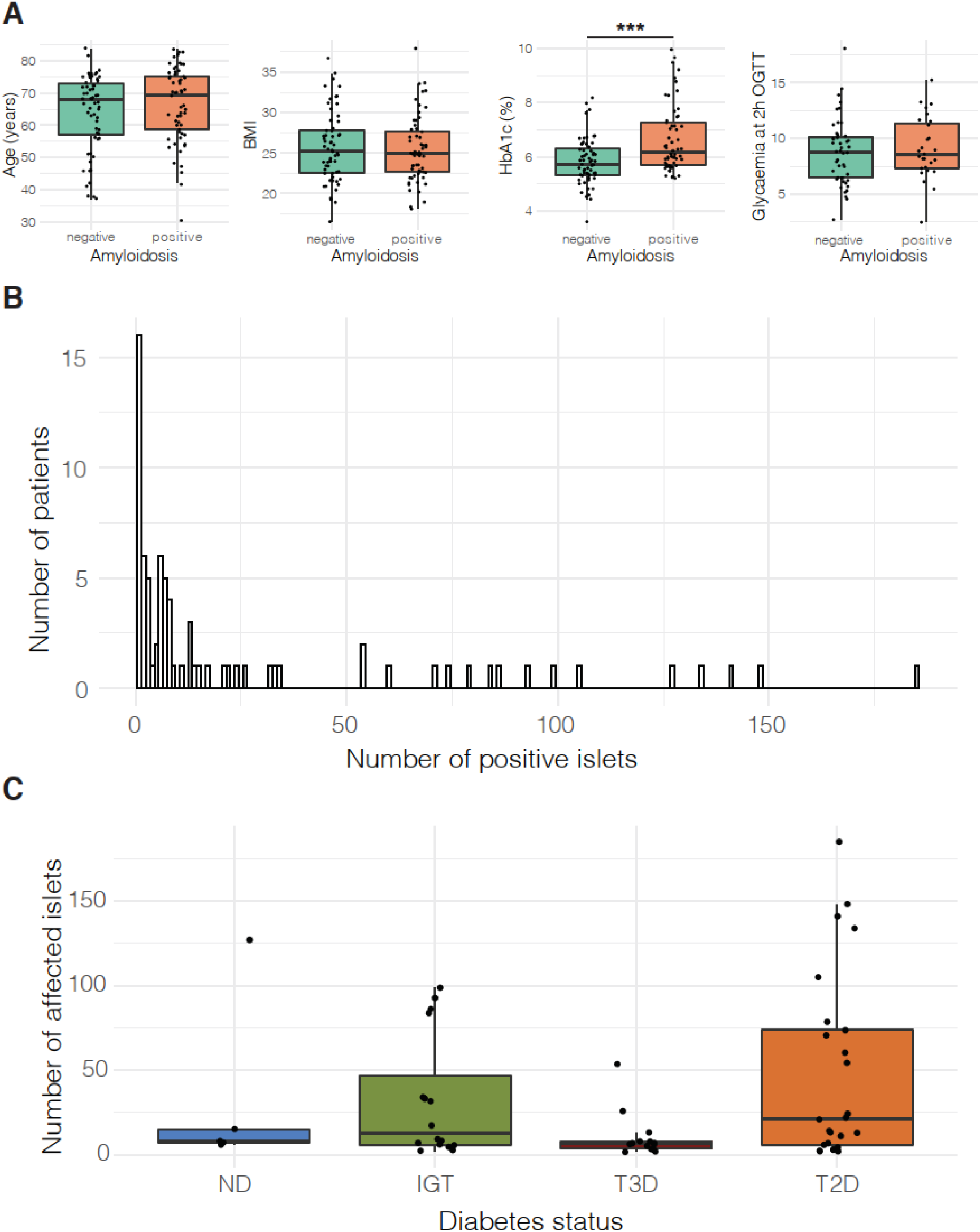
Clinical and morphological characteristics of islet amyloidosis. **(A)** Boxplots of representative clinical parameters in relation to amyloidosis positivity. Boxplot corresponds to the span between the 25^th^ and 75^th^ percentile, with horizontal line marking the median. The whiskers reach to the most extreme data point that is no more than 1.5 times the length of the box away from the box. \*\*\**p*<0.001 **(B)** Bar chart of the distribution of extent of islet amyloidosis in the cohort. **(C)** Boxplot of the absolute number of affected islets in subjects with islet amyloidosis. Boxplot corresponds to the span between the 25^th^ and 75^th^ percentile, with horizontal line marking the median. The whiskers reach to the most extreme data point that is no more than 1.5 times the length of the box away from the box.

Although a metric of limited value due to the nature and the scope of examined pancreatic islets, the distribution of the absolute number of amyloidosis affected islets pointed to a heavily left-skewed distribution. Most patients exhibited the affection of less than 50 out of the assessed minimally 200 islets and only few showing a broadly reaching islet amyloidosis (Figure 3B). The same parameter when observed in the two diabetes groups pointed out the mentioned distribution and further illustrated the presumed etiological, and possibly perduration, difference between the metabolic disorder of IGT (persistent) and T3cD (acute or sudden) (Figure 3C). The ND patient with an extensive islet amyloidosis could likely be considered an outlier.

Batch-corrected and normalized RNA sequencing data was used for data exploration considering amyloidosis status as well as other clinical (diabetes status, gender) and technical characteristics (sequencing batch). The multidimensional scaling plots only revealed a distinct separation of two patient groups, which was driven primarily by gender differences in gene expression. Based on this analysis, the batch effect appeared to be sufficiently corrected for. (Figure 4A-D).

**Figure 4.**
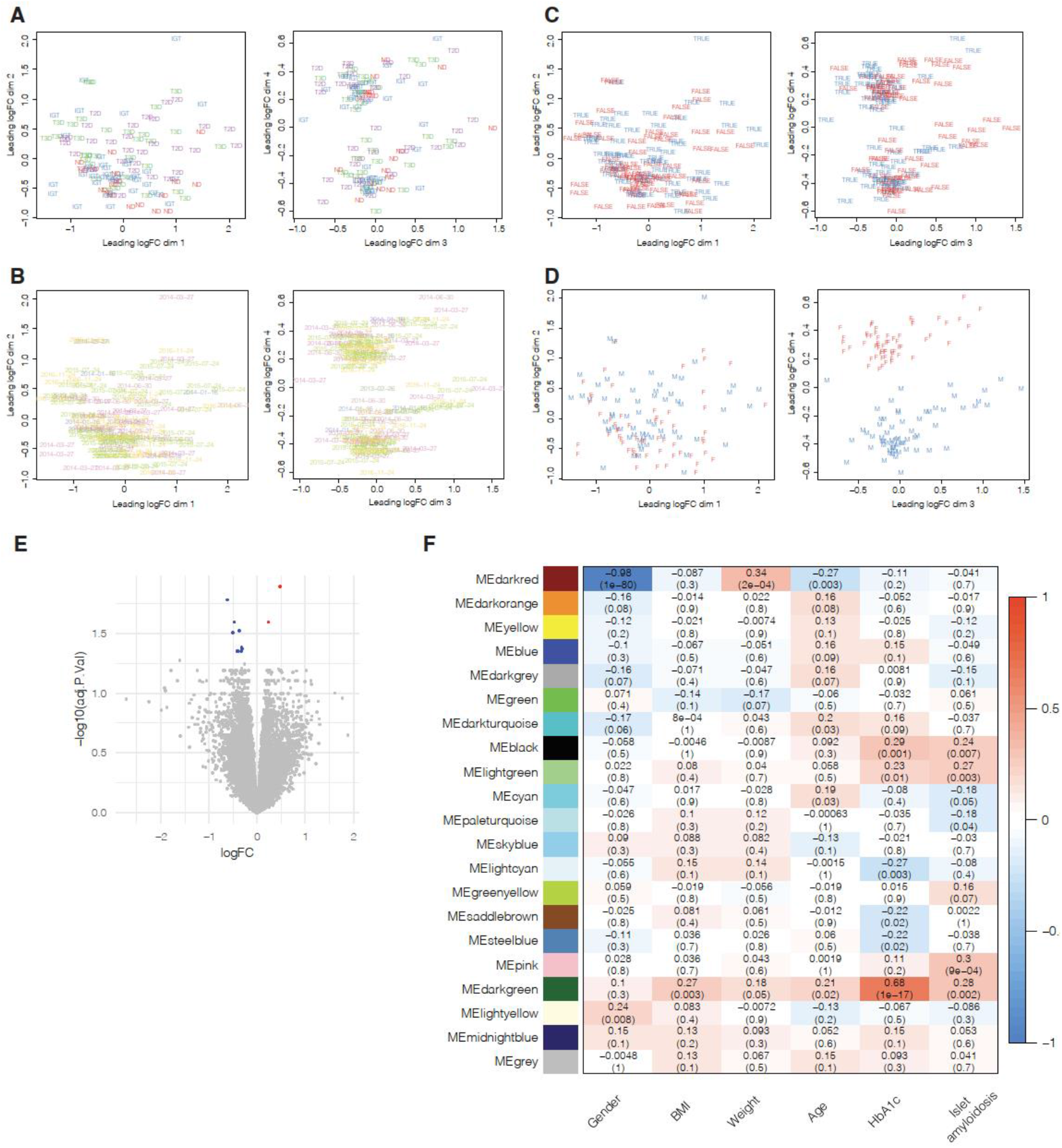
Transcriptomic signatures in islet amyloid affected subjects. **(A-D)** Multidimensional scaling plots for the first four dimensions labelled by diabetes status **(A)**, sequencing batch **(B)**, amyloid status **(C)**, and gender **(D). (E)** Volcano plot representation of differential expression analysis of amyloid positive vs amyloid negative islets. **(F)** Correlation heatmap of gene co-expression modules with relevant clinical parameters.

In an effort to pinpoint the transcriptomic changes correlated with islet amyloidosis itself rather than the metabolic disorder (prediabetes or diabetes), the information on diabetes status was included as a covariate in the linear fitting model to test for differential expression between amyloid positive or negative islets. Based on a cutoff of *p* value <0.01 and an absolute fold change >1.2, a total of 11 genes were found to be differentially expressed (Figure 4E, Table 2). Intriguingly, a long non-coding RNA was the hit with the highest fold change, while two other genes with a known relation to beta cell biology were detected as downregulated in this analysis (*PTPRN* (*p* = 0.03, logFC = -0.399) and *FURIN* (p = 0.04, logFC = -0.416)). None of these 11 differentially expressed genes had also been identified as differentially regulated in T2D donors in our previous study on a superset of the cohort presented here, supporting the validity of our approach to isolate the effects of amyloidosis from the effects of the disease/hyperglycaemic disorder.

**Table 2.** Differentially expressed genes. Islet genes differentially expressed in islet amyloidosis affected individuals. Only genes with adjusted *p* value < 0.05 and a fold change > 1.2 are reported.

| Gene.stable.ID | Gene.name | logFC | AveExpr | t | P.Value | adj.P.Val | B |
| --- | --- | --- | --- | --- | --- | --- | --- |
| ENSG0000005435 |  |  | 8,2309211 | - |  | 0,0445170 | 2,2360934 |
| 6 | PTPRN | -0,399466 | 5 | 4,3992194 | 2,40E-05 | 2 | 2 |
| ENSG0000005511 |  |  | 4,8611198 |  |  | 0,0414250 | 2,7976908 |
| 8 | KCNH2 | -0,324531 | 7 | -4,513458 | 1,53E-05 | 3 | 3 |
| ENSG0000007437 |  | - | 7,2470098 | - |  | 0,0445170 | 2,3155388 |
| 0 | ATP2A3 | 0,4028669 | 6 | 4,4147101 | 2,26E-05 | 2 | 7 |
| ENSG0000011492 |  | - | 2,8537569 | - |  | 0,0253786 | 4,0007556 |
| 3 | SLC4A3 | 0,4747201 | 5 | 4,8236809 | 4,27E-06 | 2 | 6 |
| ENSG0000013346 |  | - | 4,3588284 | - |  | 0,0424235 | 2,6801206 |
| 0 | SLC2A11 | 0,3089391 | 3 | 4,4741815 | 1,79E-05 | 2 | 8 |
| ENSG0000013592 |  | - | 4,3495235 | - |  | 0,0445170 | 2,2620230 |
| 4 | DNAJB2 | 0,3269333 | 6 | 4,3597096 | 2,81E-05 | 2 | 1 |
| ENSG0000014056 |  | - | 4,6788158 | - |  | 0,0445170 |  |
| 4 | FURIN | 0,4173172 | 6 | 4,3667022 | 2,73E-05 | 2 | 2,2676262 |
| ENSG0000018146 |  | 0,4792213 | 5,5234012 | 5,2546606 |  |  | 5,7074949 |
| 7 | RAP2B | 4 | 4 | 8 | 6,72E-07 | 0,0127772 | 6 |
| ENSG0000019622 |  | - | 3,9030119 | - |  | 0,0298799 | 3,4646294 |
| 0 | SRGAP3 | 0,3743544 | 2 | 4,6767196 | 7,86E-06 | 1 | 8 |
| ENSG0000019655 |  | - | 2,9899396 | - |  | 0,0311309 | 3,2605543 |
| 7 | CACNA1H | 0,5075699 | 5 | 4,6221361 | 9,82E-06 | 2 | 3 |
| ENSG0000027060 | AL353622. | - | 2,3856817 | - |  | 0,0164767 | 4,7494327 |
| 5 | 1 | 0,6200652 | 9 | 5,0365814 | 1,73E-06 | 3 | 3 |

To gain insights into the observed transcriptional changes, a gene set enrichment analysis of KEGG pathways was performed on the differentially expressed gene list ranked by adjusted *p* value and results were filtered using cut-offs for normalized enrichment score of 1.5 and for adjusted *p* value of 0.01. Oxidative phosphorylation, RNA transport and cytoskeleton related terms were enriched in amyloid positive islets, similarly to our previous study of transcriptomic changes in islets of T2D (Table 3). The KEGG term oxidative phosphorylation was not only interesting for its role in the islet cell energy flux, but also for a significant overlap with the terms Parkinson’s disease, Alzheimer’s disease and Huntington’s disease.

**Table 3.**
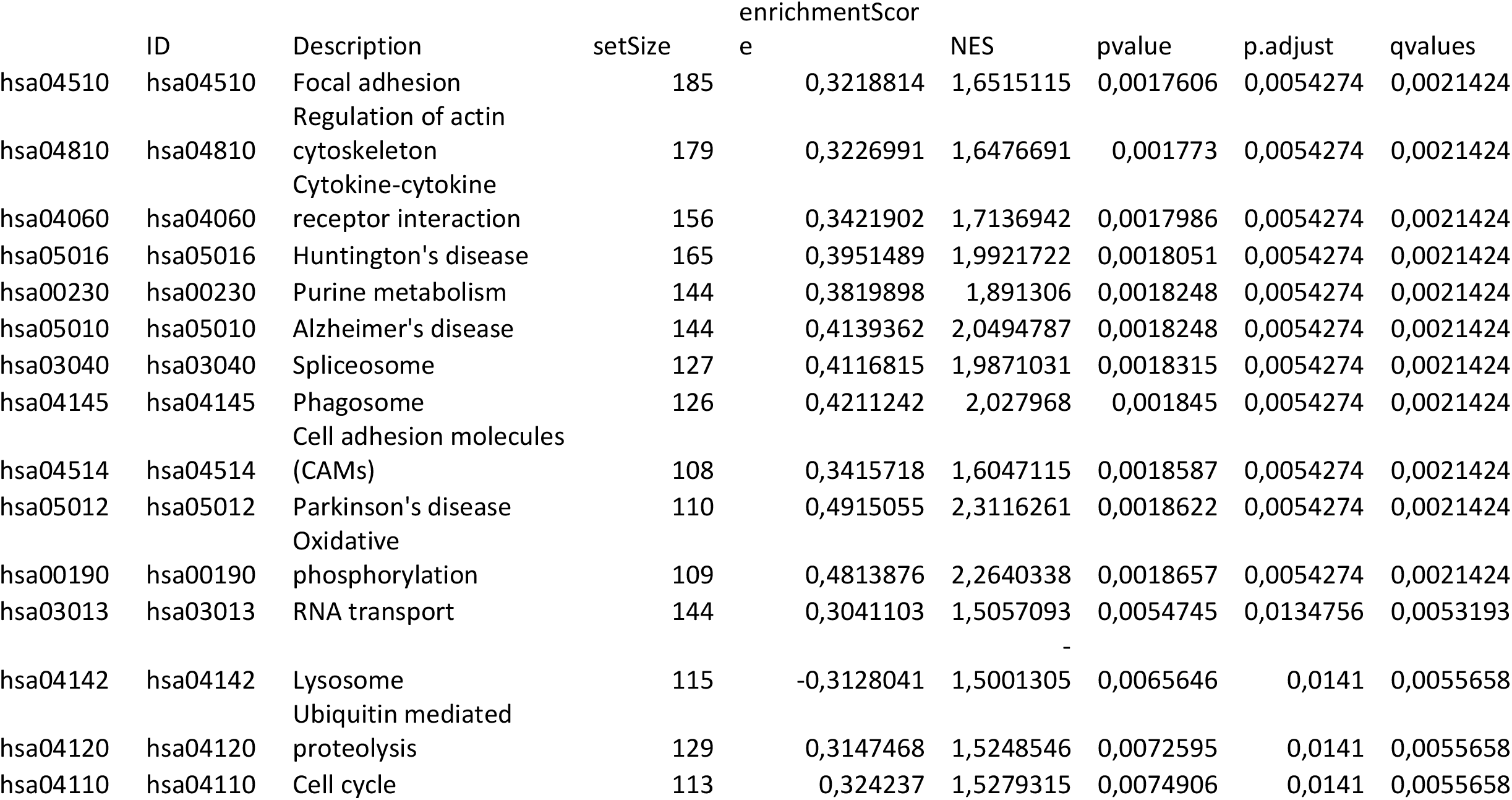
Gene set enrichment analysis - KEGG. Enriched pathways from the Kyoto Encyclopedia of Genes and Genomes in the differential expression analysis.

Testing for higher ranking of genes belonging to The Broad Institute Molecular Signatures Database v7.4 hallmark gene sets returned 15 hallmark gene sets overrepresented in amyloid positive islets with a false discovery rate of 0.05 (Table 4). Interestingly, and partially at contrast with the analysis of the KEGG gene sets, this analysis identified several gene sets primarily related to inflammation and immune response. Still, the oxidative phosphorylation gene set was also identified in this analysis.

**Table 4.**
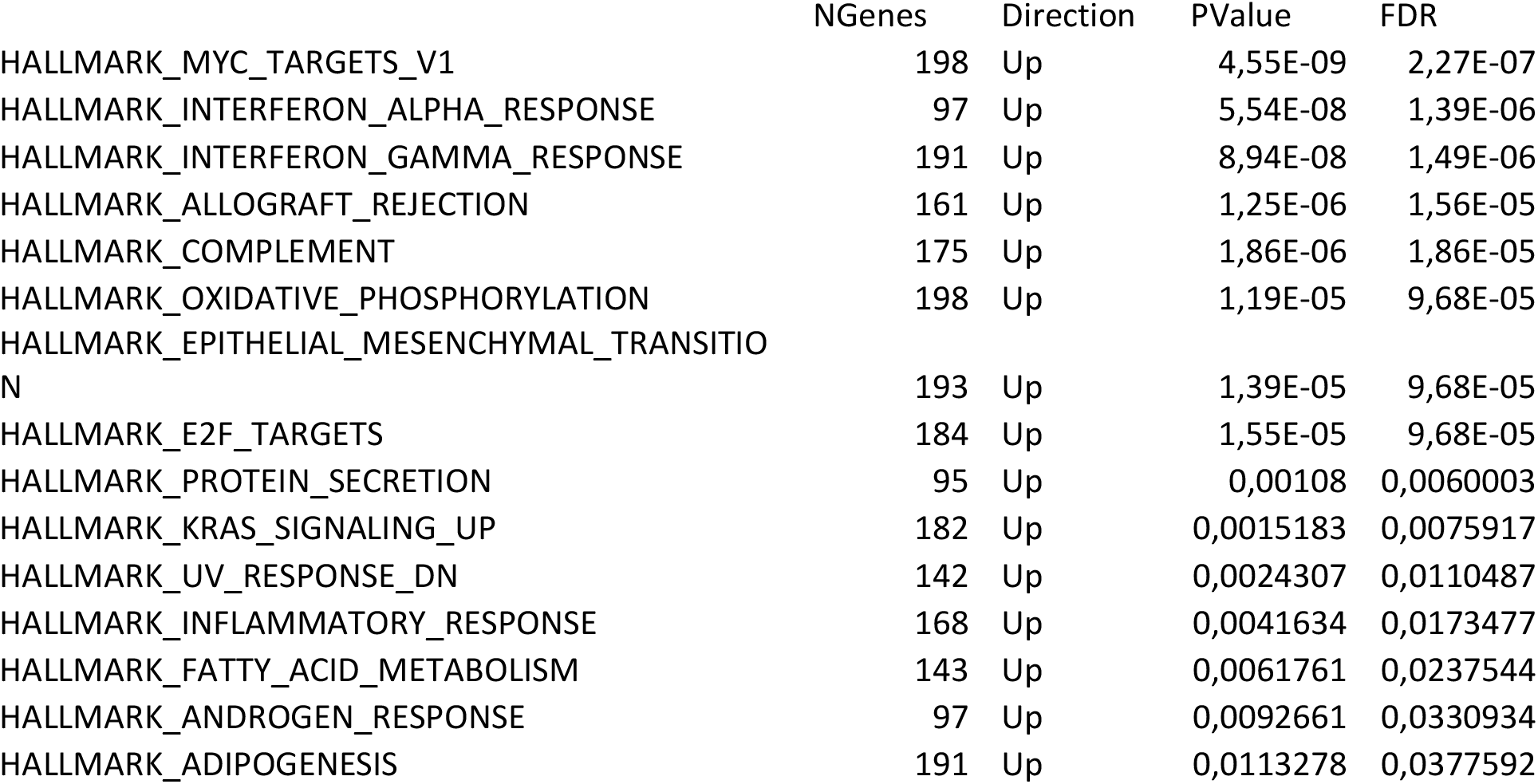
Gene set enrichment analysis - Hallmark. Enriched terms from the Molecular Signatures Database Hallmark dataset.

Exploring the transcriptomic data in pursuit of co-expressed genes with correlation to islet amyloidosis, a WGCNA was performed (Figure 4F). The pink module was identified with a weak, although significant correlation to islet amyloidosis (*r* = 0.3), which included 20 genes recognized as hub genes for their strong correlation to the module eigengene. Table 5 lists these genes, out of which four were previously reported in relation to T2D and pancreatic beta cells: ARDB1, FXYD3, F13A1 and FXYD2 (8–11). None of the module hub genes, including the four mentioned ones, have been previously associated with islet amyloidosis.

**Table 5.**
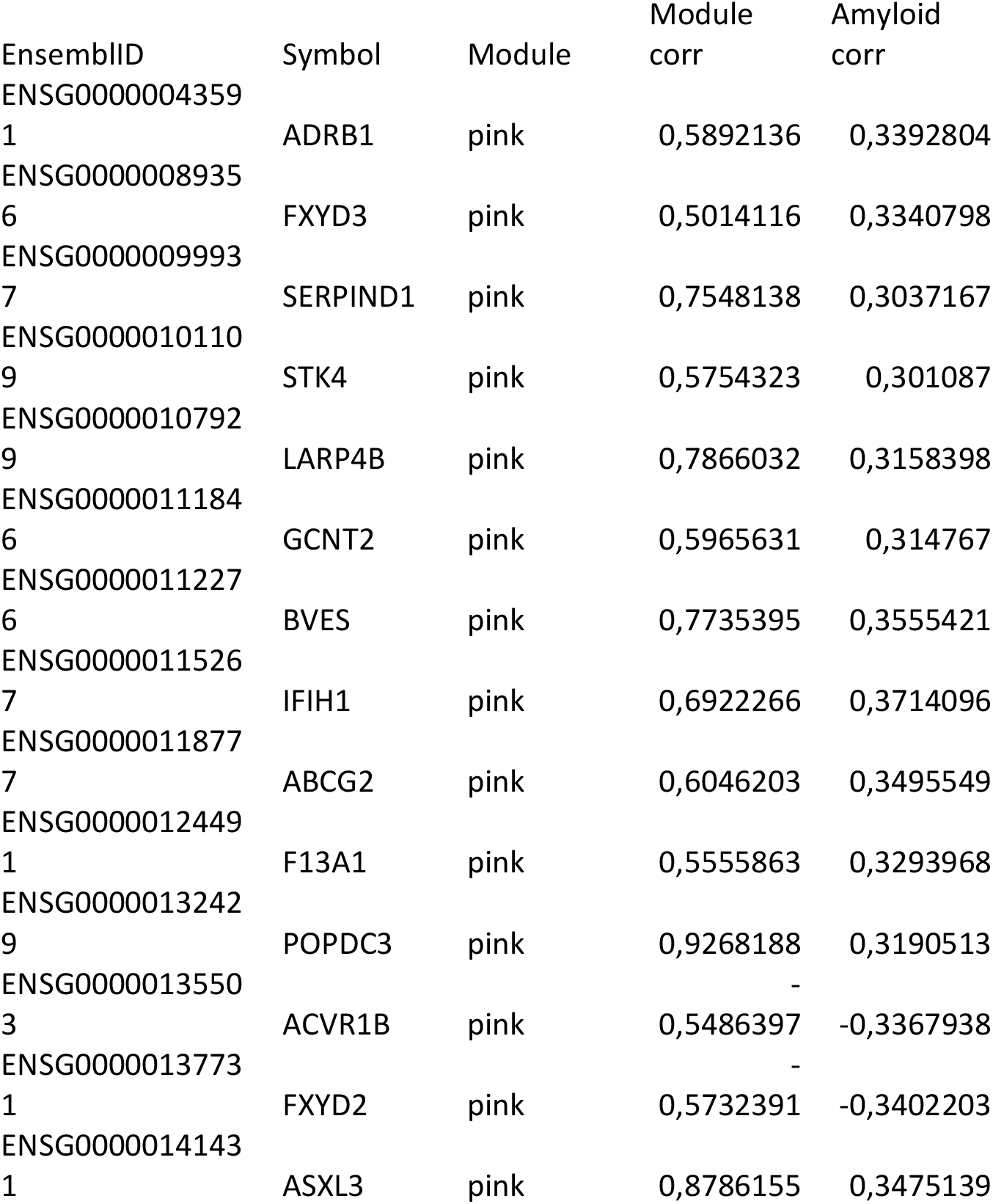
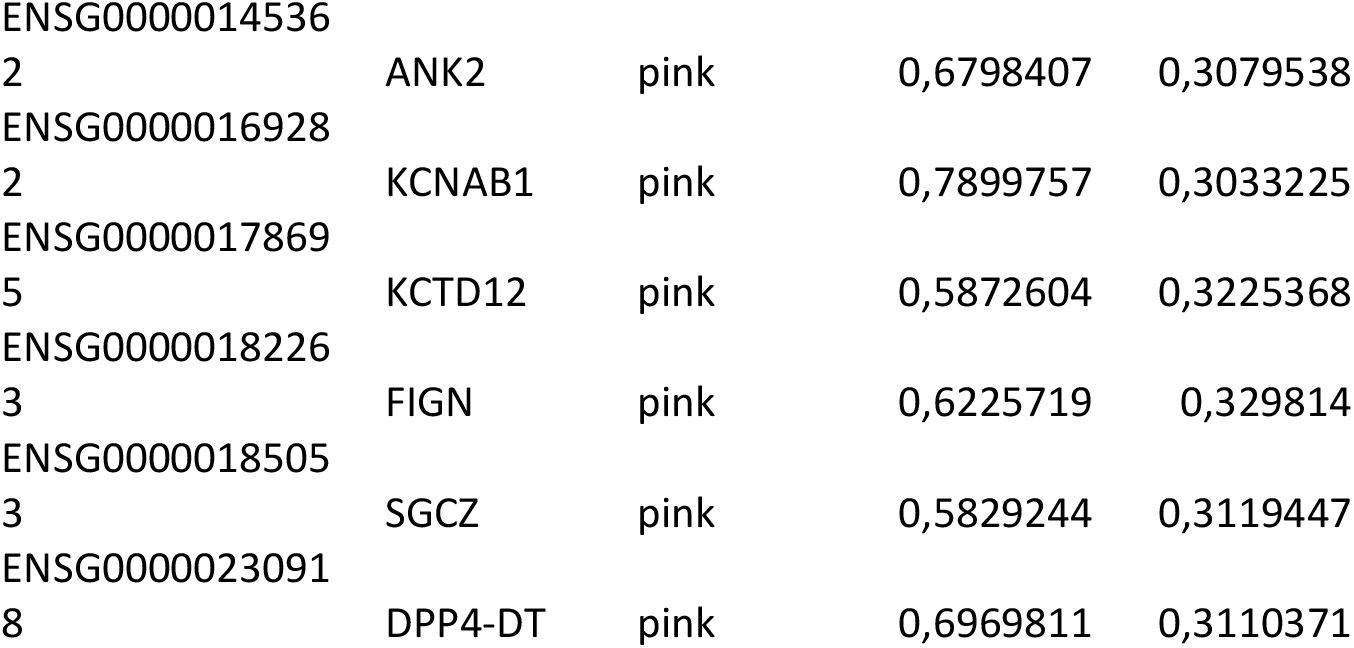
Weighted gene correlation network analysis. Hub genes of the gene co-expression module with the strongest correlation with islet amyloidosis.

## Discussion

The depth and extent of the data collected in the LIDOPACO studies was a prerequisite for this analysis, unique in its pursuit to uncover transcriptomic signatures correlated with islet amyloidosis. These investigations demonstrate how technologies relatively recently introduced into scientific practice can be utilized for extracting new knowledge regarding phenomena that are well known in the broader medical science field since many decades.

Few studies exist that directly compared the sensitivity of the two most commonly used techniques for detection of amyloid plaques - Thioflavin S and Congo Red staining.(12) The data we gathered in this study unequivocally show the superior sensitivity of Thioflavin S staining for detection of islet amyloid plaques - a procedure that is routinely used for this purpose in pathology. We hypothesize that this effect is largely conveyed by a higher contrast ratio in fluorescence-based imaging, enabling easier detection of small deposits of amyloid material. Enhanced contrast also facilitates the automated microscopy detection of signals in islet regions of interest. While Thioflavin S staining is highly sensitive for the detection of islet amyloid plaques, it comes with the cost of a loss in specificity, for which we tried to compensate by using a more stringent detection threshold.

There are several historical reports of islet amyloidosis also affecting non-diabetic individuals with varying records of its prevalence.(1,2,13–16) While the differences among the previously published investigations and also with the present study could have a biological background based in the cohort characteristics, we argue that this variability most likely stems from non-standardized methodologies for amyloid detection, as illustrated above.

Our data on the clinicopathological aspects of islet amyloidosis aligns with previous knowledge and support the notion that this alteration is positively correlated with the progression of hyperglycemia and T2D. To date there is no other detailed human clinical data available that is presented in view of pancreatic islet. The extent of islet amyloidosis among LIDOPACO donors with IGT and T3cD as presented in Figure 3C is in line with the presumed different temporal dynamics of these two entities, i.e. a chronic but slowly progressive versus an acute, but more severe deterioration of glucose homeostasis and islet function.

The unsupervised exploratory data analysis revealed no clear grouping of samples along the amyloidosis positivity line along with an incidental finding of a remarkable clustering according to gender. The following differential expression analysis was performed in accordance to the primary aim - pursuit of transcriptomic demarcations of islet amyloidosis. This entailed a complex linear modeling with multiple covariates to ensure the highest possible degree of specificity, this time with a potential sacrifice in sensitivity. Endorsing the specificity of these differentially expressed genes is the fact that none of them has been identified in the differential expression analyses on the superset of the cohort used in this study we previously published.(7) The list of differentially expressed genes, albeit short, provided valuable cues in relation to existing knowledge and for future endeavors. Interestingly, *PTPRN* (*ICA512*) and *FURIN*, genes closely associated with vesicle mediated exocytosis, were found to be downregulated in islets of subjects with islet amyloidosis. The case of PTPRN (ICA512) is especially intriguing given the high propensity of the insulin secretory granule luminal N-terminal RESP18 homology domain (ICA512-RESP18HD), to aggregate at neutral pH, i.e. in the condition encountered by granule cargoes upon their release in the extracellular milieu.(17,18) This remarkable feature sets ICA512-RESP18HD apart from other granule released peptides, which are instead soluble at neutral pH, while forming condensates at pH ≥ 6.3, as in the lumen of the trans-Golgi network and the secretory granules. Notably, ICA512-RESP18HD contains information for the sorting of peptides into insulin secretory granules and it is cleaved from the rest of the PTPRN/ICA512 ectodomain by proprotein convertases, such as FURIN. Furthermore, our data indicates that secretion of ICA512-RESP18HD from glucose-stimulated insulinoma cells is coupled with the cleavage of its C-terminal intrinsic disordered domain, which accounts for its peculiar condensing properties. In the future it would interesting to investigate whether one or both these cleavages of ICA512-RESP18HD are perturbed in conditions correlated with the formation of islet amyloidosis. Higher level analyses primarily through Gene Set Enrichment Analysis revealed some interesting patterns. Protein misfolding/ER stress signatures align with previously published studies in transgenic mice (19), but are challenged by other evidence in the same experimental model.(20) Related enriched terms referring to Alzheimer’s, Parkinson’s and Huntington’s disease are of high interest given that pathological protein aggregation is a common denominator for both amyloidosis and these neurodegenerative diseases. In addition, patients with T2D are at a higher risk to develop Alzheimer’s and/or Parkinson’s disease.(21,22) This is further supported by recent evidence of direct interaction of amyloid ß and IAPP *in vitro*.(23)

Inflammation and cytokine signaling related signatures found both in KEGG and Hallmark functional enrichment analyses do find support in the literature (24), despite the disputed importance of these processes in the pathogenesis of T2D. Our data agrees also with other conclusions of that specific study, with special emphasis on mitochondrial function and protein synthesis.

Finally, it must be noted that despite our best efforts, the extensive effects of T2D on islet cell biology could not be entirely eliminated. This limitation is revealed by the overlap with our previous findings where T2D was the focus, especially in the domain of integrative and overview analyses.(7) Nevertheless, the gene co-expression module correlating with islet amyloidosis identified in this paper might be a meaningful source of inspiration for potential future research.

## Acknowledgements

We would like to thank Juha Torkko from Solimena lab, Mario Ermacora (Universidad Nacional de Quilmes, Argentina) and Steven Kahn (VA Puget Sound Health Care System and University of Washington, Seattle, WA, USA) for valuable insights, suggestions and discussion, and Katja Pfriem for administrative assistance.

## Author contributions

JW and MD patient recruitment, surgery, provision of clinical data; ES, NK and DF tissue collection, processing and data collection; AF, DA, GB pathology and assessment of congo red staining; MB ES and NK patient data management and selection; KS immunofluorescent staining; MB development and application of amyloidosis quantification approach; FM and MB bioinformatic analysis; AS and MS conceptual insights and design, MB and MS writing of the manuscript; all authors have revised the manuscript and approved its final version.

## References

1. Opie EL. On the relation of chronic interstitial pancreatitis to the islands of Langerhans and to diabetes mellitus. J Exp Med [Internet]. 1901 Jan 15 [cited 2022 Sep 15];5(4):397– 428. Available from: https://pubmed.ncbi.nlm.nih.gov/19866952/

2. Westermark P. Quantitative studies on amyloid in the islets of Langerhans. Ups J Med Sci [Internet]. 1972 [cited 2022 Sep 9];77(2):91–4. Available from: https://pubmed.ncbi.nlm.nih.gov/4116019/

3. Paulsson JF, Andersson A, Westermark P, Westermark GT. Intracellular amyloid-like deposits contain unprocessed pro-islet amyloid polypeptide (proIAPP) in beta cells of transgenic mice overexpressing the gene for human IAPP and transplanted human islets. Diabetologia [Internet]. 2006 Jun [cited 2022 Sep 15];49(6):1237–46. Available from: https://pubmed.ncbi.nlm.nih.gov/16570161/

4. Wang F, Hull RL, Vidal J, Cnop M, Kahn SE. Islet Amyloid Develops Diffusely Throughout the Pancreas Before Becoming Severe and Replacing Endocrine Cells. Diabetes [Internet]. 2001 Nov 1 [cited 2021 Aug 3];50(11):2514–20. Available from: https://diabetes.diabetesjournals.org/content/50/11/2514

5. Marselli L, Sgroi DC, Bonner-Weir S, Weir GC. Laser Capture Microdissection of Human Pancreatic β-Cells and RNA Preparation for Gene Expression Profiling. In: Methods in molecular biology (Clifton, NJ) [Internet]. 2009 [cited 2019 Mar 11]. p. 87–98. Available from: http://www.ncbi.nlm.nih.gov/pubmed/19504246

6. Solimena M, Schulte AM, Marselli L, Ehehalt F, Richter D, Kleeberg M, et al. Systems biology of the IMIDIA biobank from organ donors and pancreatectomised patients defines a novel transcriptomic signature of islets from individuals with type 2 diabetes. Diabetologia. 2018;61(3):641–57.

7. Wigger L, Barovic M, Brunner AD, Marzetta F, Schöniger E, Mehl F, et al. Multi-omics profiling of living human pancreatic islet donors reveals heterogeneous beta cell trajectories towards type 2 diabetes. Nat Metab [Internet]. 2021 Jul 1 [cited 2022 Sep 15];3(7):1017–31. Available from: https://pubmed.ncbi.nlm.nih.gov/34183850/

8. Wieclawek A, Slawska H, Mazurek U. ADRB1 as a Potential Target for Gene Therapy of Pregnancy Induced Hypertension and Gestational Diabetes Mellitus. http://dx.doi.org/103109/106419632010532265 [Internet]. 2011 Oct [cited 2022 Nov 4];33(6):422–6. Available from: https://www.tandfonline.com/doi/abs/10.3109/10641963.2010.532265

9. Vallois D, Niederhäuser G, Ibberson M, Nagaray V, Marselli L, Marchetti P, et al. Gluco-Incretins Regulate Beta-Cell Glucose Competence by Epigenetic Silencing of Fxyd3 Expression. PLoS One [Internet]. 2014 Jul 24 [cited 2022 Nov 4];9(7):e103277. Available from: https://journals.plos.org/plosone/article?id=10.1371/journal.pone.0103277

10. Sharma PR, Mackey AJ, Dejene EA, Ramadan JW, Langefeld CD, Palmer ND, et al. An islet-targeted genome-wide association scan identifies novel genes implicated in cytokine-mediated islet stress in type 2 diabetes. Endocrinol (United States) [Internet]. 2015 Sep 1 [cited 2022 Nov 4];156(9):3147–56. Available from: /pmc/articles/PMC4541617/

11. Ferrè S, De Baaij JHF, Ferreira P, Germann R, De Klerk JBC, Lavrijsen M, et al. Mutations in PCBD1 cause hypomagnesemia and renal magnesium wasting. J Am Soc Nephrol [Internet]. 2014 Mar [cited 2022 Nov 4];25(3):574–86. Available from: https://pubmed.ncbi.nlm.nih.gov/24204001/

12. Kelényi G. Thioflavin S fluorescent and Congo red anisotropic stainings in the histologic demonstration of amyloid. Acta Neuropathol [Internet]. 1967 Dec [cited 2022 Sep 9];7(4):336–48. Available from: https://pubmed.ncbi.nlm.nih.gov/4166287/

13. Schneider HM, Stoerkel S, Will W. Das Amyloid der Langerhansschen Inseln und seine Beziehung zum Diabetes mellitus*1. DMW - Dtsch Medizinische Wochenschrift [Internet]. 1980 [cited 2022 Sep 15];105(33):1143–7. Available from: http://www.thieme-connect.de/DOI/DOI?10.1055/s-2008-1070828

14. Bell ET. Hyalinization of the Islets of Langerhans in Diabetes Mellitus. Diabetes [Internet]. 1952 Sep 1 [cited 2021 Aug 3];1(5):341–4. Available from: https://diabetes.diabetesjournals.org/content/1/5/341

15. Bell ET. Hyalinization of the Islets of Langerhans in Nondiabetic Individuals. Am J Pathol [Internet]. 1959 Jul [cited 2021 Aug 3];35(4):801. Available from: https://www.ncbi.nlm.nih.gov/pmc/articles/PMC1934824/

16. Zhao HL, Lai FMM, Tong PCY, Zhong DR, Yang D, Tomlinson B, et al. Prevalence and Clinicopathological Characteristics of Islet Amyloid in Chinese Patients With Type 2 Diabetes. Diabetes [Internet]. 2003 Nov 1 [cited 2022 Sep 15];52(11):2759–66. Available from: https://diabetesjournals.org/diabetes/article/52/11/2759/12709/Prevalence-and-Clinicopathological-Characteristics

17. Sosa L, Torkko JM, Primo ME, Llovera RE, Toledo PL, Rios AS, et al. Biochemical, biophysical, and functional properties of ICA512/IA-2 RESP18 homology domain. Biochim Biophys Acta - Proteins Proteomics. 2016 May 1;1864(5):511–22.

18. Toledo PL, Torkko JM, Müller A, Wegbrod C, Sönmez A, Solimena M, et al. ICA512 RESP18 homology domain is a protein-condensing factor and insulin fibrillation inhibitor. J Biol Chem [Internet]. 2019 May 24 [cited 2022 Sep 15];294(21):8564–76. Available from: https://pubmed.ncbi.nlm.nih.gov/30979722/

19. Huang C, Lin C, Haataja L, Gurlo T, Butler AE, Rizza RA, et al. High Expression Rates of Human Islet Amyloid of Humans With Type 2 but Not Type 1 Diabetes. Diabetes. 2016;56(August 2007).

20. RL H, S Z, J U, K A-M, SL S, SE K. Amyloid formation in human IAPP transgenic mouse islets and pancreas, and human pancreas, is not associated with endoplasmic reticulum stress. Diabetologia [Internet]. 2009 Jun [cited 2021 Aug 3];52(6):1102–11. Available from: https://pubmed.ncbi.nlm.nih.gov/19352619/

21. Cheong JLY, de Pablo-Fernandez E, Foltynie T, Noyce AJ. The Association Between Type 2 Diabetes Mellitus and Parkinson’s Disease. J Parkinsons Dis. 2020 Jan 1;10(3):775–89.

22. Cooper C, Sommerlad A, Lyketsos CG, Livingston G. Modifiable predictors of dementia in mild cognitive impairment: a systematic review and meta-analysis. Am J Psychiatry [Internet]. 2015 Apr 1 [cited 2022 Sep 15];172(4):323–34. Available from: https://pubmed.ncbi.nlm.nih.gov/25698435/

23. Wang Y, Westermark GT. The Amyloid Forming Peptides Islet Amyloid Polypeptide and Amyloid β Interact at the Molecular Level. Int J Mol Sci [Internet]. 2021 Oct 2 [cited 2022 Sep 15];22(20). Available from: https://pubmed.ncbi.nlm.nih.gov/34681811/

24. Hogan MF, Ziemann M, N HK, Rodriguez H, Kaspi A, Esser N, et al. RNA-seq-based identification of Star upregulation by islet amyloid formation. Protein Eng Des Sel [Internet]. 2019 Feb 1 [cited 2021 Aug 3];32(2):67. Available from: /pmc/articles/PMC6908819/

